# Intradisciplinary Growth of Sustainability-Minded Engineers through Conservation Technology

**DOI:** 10.1101/2023.07.03.546429

**Authors:** Andrew Schulz, Cassie Shriver, Anika Patka, Caroline Greiner, Benjamin Seleb, Rebecca Watts Hull, Carol Subiño Sullivan, Julia M. Sonnenberg-Klein, Roxanne Moore

## Abstract

**Background:** The need for sustainability-minded engineers prepared to address complex societal challenges has grown exponentially in recent years. Frameworks like the United Nations (UN) Sustainable Development Goals (SDGs) have begun to drive structural changes in engineering education, including new ABET accreditation focused on sustainability. The new field of conservation technology allows engineers to develop sustainability competencies and identities as conservationists and environmentalists.

**Purpose:** This manuscript describes an assessment of student identity development in conservation and environmentalism in the GaTech4Wildlife Vertically Integrated Project (VIP) course at Georgia Tech. The course uses the principles of Human-Centered Design along with the UN Sustainable Development Goals and project-based learning to solve conservation-oriented, real-world problems and develop sustainability-minded engineers.

**Design/Method:** Undergraduate students participated in the course and utilizing both in-person interviews and post-course assessment, students were assessed for course themes and identities. The sample consisted of students from the College of Engineering and the College of Computing.

**Results:** Since 2019, over 50 students have participated in this Tech4Wildlife course. Based on surveys and interviews of nearly 20 of the most recent students, students transitioned from identifying as engineers and coders with no sustainability knowledge to nearly doubling their identity measures as conservationists and environmentalists after only one semester.

**Conclusions:** To teach the next generation of sustainability-minded engineers, interdisciplinary, project-based courses grounded in Sustainable Development Goals may offer a meaningful pathway for students to develop both technical skills and conservationist identities.

## Introduction

In 2015, the United Nations released the Sustainable Development Goals (SDGs) to provide a framework for addressing global challenges across non-profits, industry, politics, and education [1]. The vision of a sustainable future on earth was condensed to 17 individual goals that each provide far-reaching research applications for higher education, similar to those of the National Science Foundation’s (NSF) Big 10 Ideas and the Grand Challenges proposed by the National Academy of Engineering. Many colleges and universities use the SDGs to guide research, scholarship, and service [2]. The SDGs have also recently gained pedagogical traction with educators as High Impact Practices, or HIPs, guides for classroom learning settings [1, 3, 4, 5]. Two HIPs are capstone projects and community-based learning[6]. Curricular integration of the SDGs in higher education programs varies widely from “add-on” approaches that create a new major or certificate to integration across the core curricula and even profound curricular transformation described as “Education as Sustainability” [7]. A primary driver of the SDGs is the need to develop new and innovative solutions to sustainability challenges caused by climate change, food insecurity, public health crises, and inequality[8].

In 2001, the Vertically Integrated Projects (VIP) model was developed at Purdue University, enabling faculty to embed large student teams in their research. The model has been implemented at more than 40 institutions throughout the globe focused on the following core tenets: 1) projects are embedded in faculty mentor’s research, scholarship, or exploration, 2) projects are long-term and large-scale, lasting many years, 3) the program is curricular, 4) students can participate for multiple semesters; 4) learning outcomes include both disciplinary and professional skills [9]. While not required, multidisciplinarity is a hallmark of VIP Programs because large-scale problems are multidisciplinary by nature. Through VIP, faculty can develop new ideas (such as conservation technology) in a low-stakes setting and engage undergraduates in novel learning experiences without significant changes to the curricula or requirements of typical engineering degrees [10].

One of the unique attributes of sustainability solutions and the VIP program is that many of the solutions address a diverse set of fields. Currently, engineering education has many initiatives that work on educating interdisciplinary teams, including Capstone Design Teams [11], Freshman course experience [12], and interdisciplinary research projects [13]. However, one gap in these education systems is that many of these interdisciplinary fields have mono-disciplinary solutions. Fields such as bio-inspired design often address engineering challenges using structure-function mechanisms to develop materials[14, 15, 16, 17], robotics [18, 19, 20, 21], or algorithms [22, 23, 24]. In some of these fields, although the team might be interdisciplinary, the solutions are focused on one particular field. Conservation Technology (CT) is an overarching term for wildlife and environmental conservation developments. Much existing CT implements modern hardware and software design processes to improve ongoing conservation efforts and to initiate previously under-addressed efforts [25]. Some of the major goals of CT are to improve outdated equipment, increase accessibility to tools, and use modern technology to address conservation problems in entirely new ways. For wildlife, in particular, CT is being developed for both animals’ natural environments (in-situ) and captivity settings (ex-situ)– e.g., foxes and elephants, respectively) [26]. Initially, the conservation biology community was dismissive of new technologies, but biologists have recently begun to embrace and even assist in developing new conservation tools and solutions [27]. Unlike many mono-disciplinary engineering endeavors, every CT development must be adequately defended by establishing the necessary bridges between the conservation community, technologists, and policymakers [28, 25]. New technology is not considered conservation technology until its usefulness and success are demonstrated in the appropriate setting. In this field, there is a requirement for interdisciplinary and diverse sets of skills in the room, and when we look at the challenges, many of these are interdisciplinary as well. Many CT solutions include engineering requirements, public policy issues, wildlife management, and data analytics and ethics.

The field of CT utilizes collaborations with conservation practitioners, such as zoological organizations, which have a history of experience in sustainability education. Interdisciplinary solutions such as bio-inspiration at zoological institutions allow for applications in biodiversity conservation [29]. However, zoological organizations focus on different goals than universities [30]. This is because the primary goal of zoos is education about the species in the zoo, with animals serving as ambassadors for species in the wild. Nearly 600 million people visit over 1300 zoos and aquariums worldwide each year [31]. While education is a primary goal of zoos, it is not a chief reason that people go to zoos [32]. At zoos, aquariums, or even museums, visitors can make personal connections and control their own learning environments, which has long-term impacts (nearly 10% of visitors change their daily actions at home, such as reduction of the use of single-use plastics or discussing environmental issues with peers) [32].

Bio-inspired design is a multidisciplinary approach to solving novel engineering problems, drawing inspiration for engineered systems from biological mechanisms [33]. While a bio-inspired curriculum may build a conservation or sustainability focus as a secondary effect through inspiring connections with nature and cross-disciplinary collaboration, it does not explicitly address the need for more green and sustainability-focused education[34] (**Figure 1**A). In this paper, we hypothesize that by utilizing established zoological practices in the engineering classroom, we can create a more sustainable and environmentally conscious engineer without the need for additional environmental engineering coursework.

**Figure 1:**
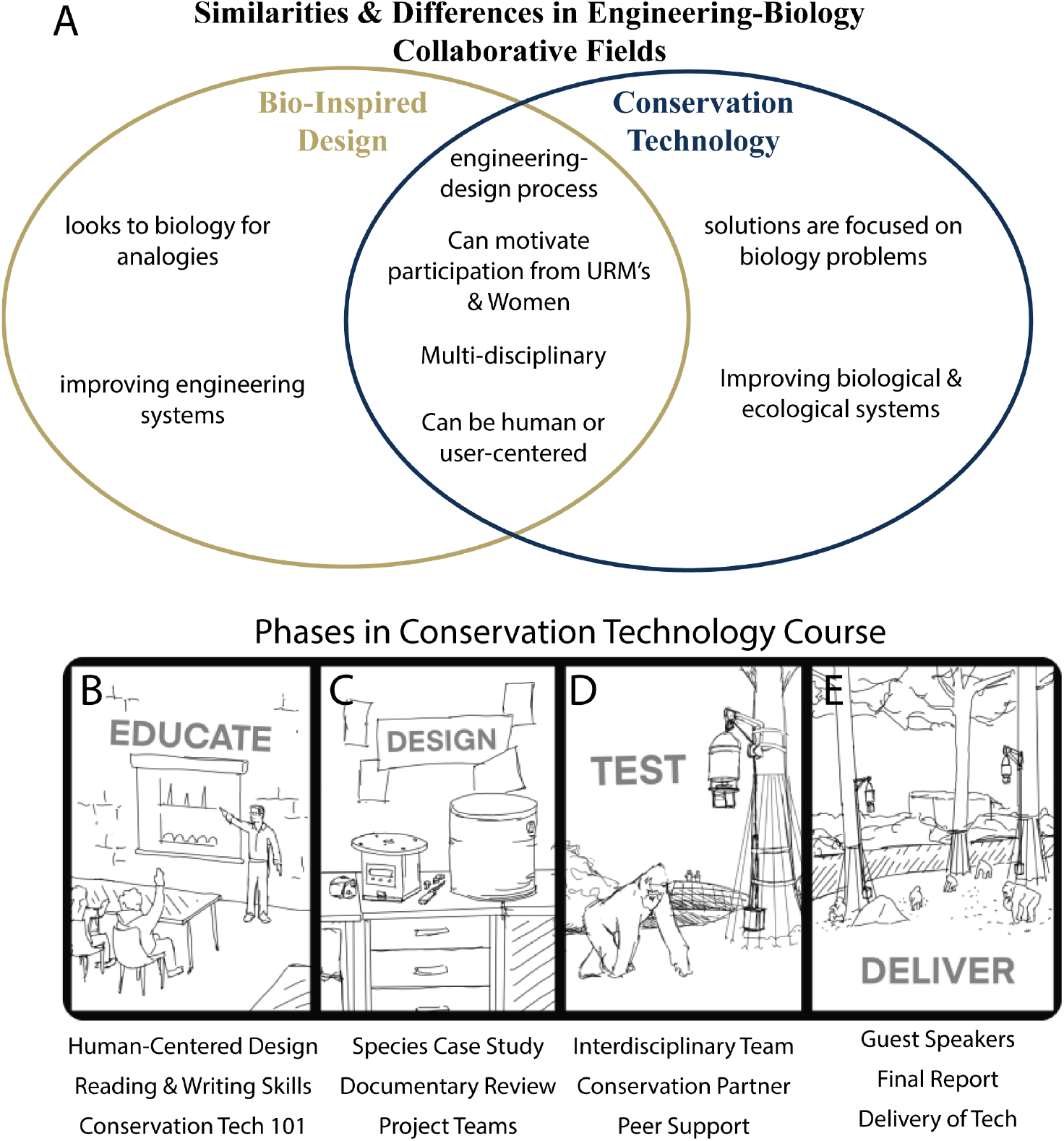
A) Similarities and Differences between Bio-Inspired Design (BID) and Conservation Technology (CT) as specific fields for Engineering-Biology educational frameworks. B-E) Phases of the conservation technology project-based course including B) Education phase of the course where students directly interface in an active-learning classroom where they are split into teams for C) Designing their specific conservation technology solutions before D) testing these solutions with their project sponsors and iterating through the engineering design process until the teams E) deliver the finalized project.

### The VIP Model

Through VIP, faculty embed large student teams in their research, scholarship, and exploration work. Teams are vertically integrated, meaning that they include sophomores (2nd year), juniors (3rd year), and seniors (4th-5th year), as well as graduate students and post-doctoral scholars. Students earn 1-2 credits per semester, with multiple semesters of participation being worth the same number of credits as a conventional class. Faculty can maintain large teams, 20 students on average because returning students take on additional leadership responsibilities[35, 36]. VIP teams are established by faculty request, and projects include funded research, areas of interest (such as service learning), and pedagogical improvements in various engineering education fields, from Mechanical Engineering to Electrical and Computer Science[37]. The Georgia Tech program was established in 2009, and in Spring 2023, it served over 80 teams and 1,700 students.

While VIP projects vary, commonalities across the program are the grading framework, the fact that teams meet weekly, and typical team structures. Students are graded in three equally weighted areas: contribution to the project, documentation, and teamwork. Contributions to the project include work that moves the project forward and work students do to get up to speed (tutorials, etc.), with differing expectations for new and returning students and students of different academic ranks. Contributions are assessed through instructor knowledge of the project, weekly check-ins, instructor-defined deliverables (presentations, formal write-ups, etc.), and evidence of contributions in the documentation. Documentation, the second graded category, includes individual documentation (VIP notebook, electronic notebook, web-based project management platform, etc.) and contributions to team-level documentation that will be used to continue the project the following semester. Teamwork, the third graded category, is assessed through peer evaluations and observations in weekly meetings. Students are given feedback at the middle and end of each semester. Mid-semester feedback is advisory, indicating the level of students’ work (“if you keep doing what you’re doing, you’ll get a B”), with suggestions for improvement.

Teams meet for 50 minutes each week (a scheduled class), and they are typically organized into subteams. At weekly meetings, students and subteams report on the work done in the previous week and identify tasks for each student/subteam for the upcoming week. In most teams, subteams work on aspects of a central project. Service-learning teams often work on multiple related projects, which enables subteams on different projects to help and learn from each other.

A variety of studies have been conducted on Georgia Tech’s VIP Program. An analysis of institute exit surveys found that compared to non-participants, VIP participants more strongly agreed that their Georgia Tech educations supported their ability to work in multidisciplinary teams, their ability to work with individuals from diverse backgrounds, and their understanding of technology applications relevant to their fields of study, with meaningful effect sizes on all three (Cohen’s *d* of .2 or greater) [38]. Social network analysis of peer evaluations was used to explore student interaction by significance and background further. Results showed that within their teams, students interacted more often with students from other majors and more often with students of other races/ethnicities. This indicates that students within teams cross disciplinary lines and that interaction is shaped by project dynamics instead of race/ethnicity[39]. Analysis of student enrollment showed that students return for second, third, and subsequent semesters at the same rate, regardless of race/ethnicity, an essential measure of equity [40]. Studies have also shown leadership growth within VIP. Cross-sectional and longitudinal analysis of peer evaluations shows that, as rated by peers, students take on increasing leadership responsibilities through their third semester of participation. That leadership is associated with experience on the team and not academic rank[35, 36].

The Tech4Wildlife VIP team was established in **Fall 2019**. The team grew out of another VIP team, **HumaniTech**, so the modules developed for Tech4Wildlife have been used since **Spring 2019**.

### Human-Wildlife-Centered Design as a Framework for Conservation Technology

The Tech4Wildlife VIP course is built around the four-phase framework of human-wildlife-centered design (**Figure 1**B-E). The semester begins with **education** on broad concepts, including human-centered design, conservation science, frugal technology, and technological interventions (**Figure 1**B). We then pair interdisciplinary student groups with conservation leaders in the field to provide specific expertise and allow them to **design** a conservation tool to help the organization and the impacted species (**Figure 1**C). Students work through the engineering design process to **test** the validity of their design (**Figure 1**D) and, after iteration, **deliver** the final conservation tech tool[41] (**Figure 1**E). More detail can be found in Schulz et al. [41].

Conservation technology learning objectives for are:[41]:

- Identify examples of conservation technology & human-centered design
- Review and critique current CT projects using human-centered design principles
- Explain the historical issues in conservation technology and become an advocate for utilizing human-centered design to help wildlife and the environment
- Analyze current CT practices, including tracking, monitoring, software, and hardware technologies used in the industry
- Design and submit an interdisciplinary CT project to an online digital maker space (https://conservationxlabs.com/digital-makerspace)
- Communicate with international leaders in species conservation The course has a range of assessments for feedback and grading. These assessments include:
- Notebook - Includes what a student did that week and what they plan on doing next week. It adheres to the VIP standards of facilitating learning and documentation.
- Annotated Bibliography - This is a beginning-of-the-semester assignment that helps students understand exactly what the class is about by examining existing solutions in the wildlife world using technology
- Peer Evaluations - An important component of the VIP team experience is peer evaluation. Students evaluate classmates with whom they work, with one evaluation midway through the term and one at the end of each semester.
- Species Case Study - This assignment guides students through understanding their target species. Each student writes a two-page reflection on a current species on the red list of the International Union for Conservation of Nature (IUCN). This assignment is given after the first few weeks of class, and students are given the agency to select their preferred deliverable, such as an infographic, essay, or recorded presentation.
- Presentations - Each team is expected to give a 10-minute presentation on their progress for their project several times during the semester. There is also a final, culminating presentation.
- Final Project Report - The final report is both a grading opportunity and a transition document for the team continuing the project next semester. This report should expand on the final presentation with additional details.

We proceed to discuss each phase of the four phases of the course.

### Education Phase - A Sustainability Foundation

The Education and Design phase encompasses the first few weeks of the course. The Education phase is the first five weeks primarily focused on active learning scenarios before students break up into their groups. Returning students to the course then have time to refresh their projects and prepare for which new members they would like to recruit to their team.

We are onboarding new students by introducing the human-centered design, which incorporates the perspectives of intended users throughout the design process to enhance product reception and success. This helps students transition to the fundamental concept of the course, *human-wildlife-centered design*, which combines human-centered design with human-wildlife conflict. In conservation biology, human-wildlife conflict is a prevalent challenge, with examples ranging from humans invading animals’ native habitats to disease transmission of urban wildlife such as coyotes, foxes, and monkeys in dense urban areas.

During onboarding, students learn methods for creating a successful conservation technology artifact. Through this process, students build a foundation for collaboration between unrelated disciplines and a framework for working with external partners.

During the first phase of the course, we have an initial phase of education for the students. The primary focus of this phase is to educate folks who are not traditionally in sustainability-focused classes with the foundational knowledge necessary to succeed. Additionally, we have returning students who help lead this discussion as well. The content and examples utilized each semester are updated based on research conservation events and challenges, so returning students do not receive the same examples but do get refreshers on the content.

In conservation biology, human-wildlife conflict is a prevalent challenge, with examples ranging from humans invading animals’ native habitats to disease transmission of urban wildlife such as coyotes, foxes, and monkeys in dense urban areas. The onboarding modules are:

- Introduction to Human-Centered Design, Conservation Technology, and the Sustainable Development Goals
- Human-Centered Design and Context of Use
- Effective Reading, Writing, and Interdisciplinary Teamwork
- Conservation Technology 101 - Biology: Ecology and Behavior
- Conservation Technology 101 - Indigenous Design Solutions

The introduction of Human-centered design is grounded in the IDEO framework discussed in the project-based engineering education setting by Mueller et al. [42]. We discuss and introduce the United Nation’s sustainable development goals framework as a prompt for later assessments discussed in the design phase. We proceed in the following weeks with a deep dive into human-centered design, interdisciplinary teamwork skills, and a foundational groundwork of conservation technology and biological sciences. The framework of biological sciences is essential as these conservation technology solutions require working with biological collaborators that are hypothesis-driven with their projects. In contrast, engineers and computer scientists traditionally focus on design artifacts and their functionality.

### Design Phase - The Conservation Tools of Tomorrow

We proceed with the design phase, where students learn potential conservation technology applications individually through a documentary review and case study. The documentary review allows students to watch a specifically selected list of documentaries and interpret potential technological solutions that could be implemented. Additionally, students can include a discussion of possible improvements to existing technologies depicted in the documentary. This completes the individual assignments portion of the course, and we proceed with team assignments where students engage with varying teams.

Once the course fundamentals and **conservation education** have been covered, we form inter-disciplinary teams of engineers, biologists, and computer scientists with various partners (**Figure 2**A-C). These teams are paired with conservation leaders for local, domestic, and international projects to establish **multinational collaborations**. For each project team, students work to bring forward indigenous design and work to counter the historical colonization of conservation that has been done in the CT space. Finally, students work to conduct or inform **ethical experiments** by working on IACUC and IRB protocols.

**Figure 2:**
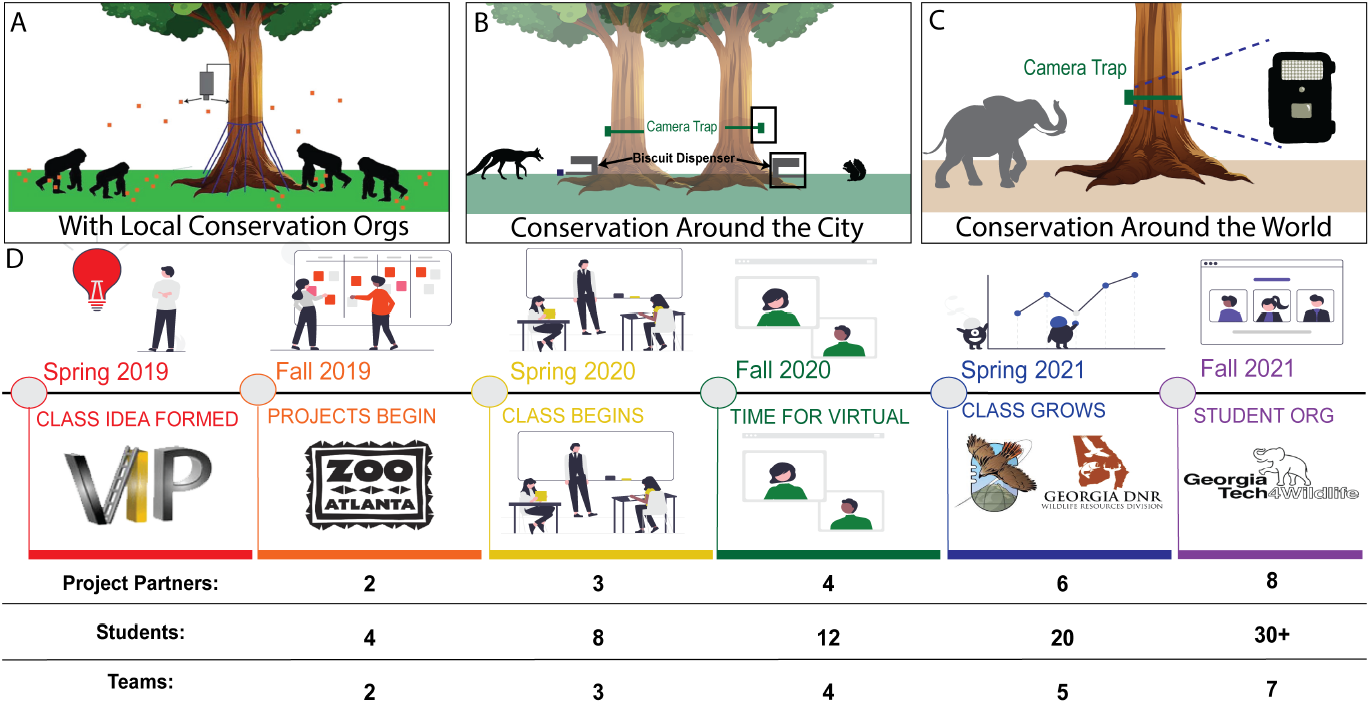
A-C) Common course project types include examples of A) working with Zoo Atlanta on an automated forage feeder[44], B) Fox biscuit dispensers for rabies vaccination, and C) Wildlife cameras to identify elephant human-wildlife conflict.

During this group project work, teams meet weekly outside the class as interdisciplinary teams and document their progress in notebooks. There is an additional check-in midway through the course with peer reviews of teammates. Instructors use peer review results to provide students with constructive feedback. The course spans several semesters, and the course alumni mentors and new members. The peer-support aspect is a cornerstone of the successful VIP program at tech[43].

### Test Phase - Engineering Iteration of Conservation Tech

After the team has been working on their project in the middle of the semester, they will proceed to the test phase. This phase begins when the students have a completed test prototype and sit down with the project partners from the conservation community. During this phase, students engage with project partners to test their devices in their chosen study area. This could be the distribution of camera traps around campus, going to the zoo to hang up a feeder, or sending frugal hardware to the project partner.

The testing phase yields noteworthy results for the student’s projects, unveiling the project prototypes’ strengths and limitations per the project partners’ advice. Through peer analysis with the project partners, students identified vital design flaws, technical glitches, and unforeseen challenges during testing, including printed parts not interfacing, material constraints due to environmental conditions, and debugging code in the field. These findings enabled students to refine their prototypes and enhanced their ability to adapt to dynamic situations and think critically when faced with unexpected obstacles. Finally, the testing phase fosters collaboration, promoting interdisciplinary discussions and knowledge exchange among students, instructors, and project partners.

This testing is iterative and likely occurs several times as students develop through different prototype models.

### Deliver Phase - Tools in the Field

During the end of the semester, we focus on connecting students with experts in the field that have or are currently working towards delivering completed and tested conservation technology projects in varying communities worldwide. This phase allows students to get conservation technology experts’ final advice on their projects before they are delivered to the project partners.

These conservation technology experts in hardware, software, scientific discovery, and frugal development all provide valuable insight to students. The experts brought into the class often do not know many of the current projects but allow students to get expert advice near the end of the semester when they are putting the final touches on their final deliverable prototype at the end of the semester. Over time the course has evolved, and the delivery phase of this project might be one part of a whole design. Therefore it is essential to know that the testing and delivery phases can often be multi-semester for the entire device for students’ delivery to the project partners.

#### Project Case Study

To provide a case study of an interdisciplinary project between engineers and scientists, we will discuss an ongoing project that began in the Spring of 2020. The external partner was Zoo Atlanta, which asked the Tech4Wildlife team to help them encourage more realistic foraging behavior for gorilla populations in captivity (ex-situ). To make captive gorillas’ diets and feeding schedules more similar to what they would experience in the wild, the team designed a randomized automated feeder. Students were initially provided a rough idea of what the project would entail based on the zoo’s identified needs. The zookeepers had unsuccessfully attempted to connect a car battery and a store-bought deer feeder but could not re-purpose the device within the Zoo Atlanta gorilla habitat. They were excited to advise a team of undergraduates in developing a better solution. The team began by meeting with the head gorilla keeper at Zoo Atlanta to understand what the device would need to do and how to test the validity of a finalized prototype.

The device’s final design was completed at the beginning of Spring 2022. Students received consistent and frequent feedback over the four semesters and underwent several iterations before achieving the final design. The final project is easily replicated using handheld tools, a 3D printer, and a mass-produced printed circuit board (PCB) design, allowing zookeepers and other conservation community members with less engineering experience to implement this design with minimal instruction. Continual communication with Zoo Atlanta’s conservation experts supported project success. The final product met all of the gorilla keepers’ needs. It provided the team of engineers, computer scientists, and biologists with invaluable hands-on experience designing solutions that address human and wildlife needs. The hardware for this case study was eventually published in the journal HardwareX by Jadali et. Al[44].

## Methods

### Qualitative Interview Data

To understand the student impacts of the popular Conservation Technology course at Georgia Tech, we sought to identify the grounding themes of the course. We performed traditional thematic analysis through formalized in-person interviews with approval from Georgia Tech’s Institutional Review Board (IRB). Past and current students in the GaTech4Wildlife course were asked to participate, and we interviewed nine individuals. Each interview lasted no longer than 30 minutes, in which we employed the following structured interview protocol:

1. If you have taken multiple VIP classes, how did the GaTech4Wildlife VIP course compare to other VIP classes you have taken?
2. What aspects of the GaTech4Wildlife VIP course did you find to be effective?
3. Are there any specific elements of the GaTech4Wildlife VIP course that built interest and engagement?
4. How did the traditional assignments (case study, documentary review) affect your experience in the GaTech4Wildlife VIP course?
5. How did the experts/guest speakers affect your experience in the GaTech4Wildlife VIP course?
6. How did the project-based learning format affect your experience in the GaTech4Wildlife VIP course?

During the interviews, participants could discuss their answers without any time limitations. Each interview was concluded by asking the participants if they had any other feedback or thoughts they would like to share regarding their experience in this course. Transcriptions were completed using Rev.com – names and personal information were not included in the interviews or transcriptions. Once transcribed, we used a grounded theory approach to develop a code book and guide a thematic analysis. Emergent themes are defined and then exemplified using illustrative quotations from students. A summary of the common themes and illustrative quotations are detailed in [45]. When the Conservation Technology course was first established at Georgia Tech in the Fall of 2019, we started with a class size of four and two projects. In the Spring of 2022, we had a class size of 20 students working on six different projects partnered with various wildlife conservation organizations partners. Based on our analysis, we found that the following four themes from our interviews all appear in over 75% of representative interviews:

- Hands-on team project experiences in conservation tech
- Exposure to research opportunities for individuals to apply learned knowledge to conservation
- Eye-opening perspective on using technology to conserve wildlife
- Connections made among varying disciplines during the course

### 0.1 Quantitative Survey Data

To further evaluate how VIP participants felt about the course, we created a 15-question, 5-minute online survey using the Qualtrics software platform. The survey was divided into three main sections: (1) acknowledgment of consent to participate in the research study, (2) background information, including gender identity (if comfortable), amount of time in VIP, and college at Georgia Tech (3) gauging pre-class and post-class identities as well as the importance of thematic analysis results to recruitment and participation in the course. The survey concluded with a question about end-of-course takeaways.

We distributed the survey to students participating in the VIP course during the 2022 and 2023 spring and fall semesters. The survey was distributed after the final reports were turned in, as there were survey questions about them. No incentives were given to survey participants, and it was purely voluntary. We had 5 participants fill out the survey from the Spring of 2022 and 49 from the Fall of 2022 semesters for a total of 18 student participants.

We used Qualtrics reports to obtain distributions of answers, averages, and standard deviations, and then RStudio and Adobe Illustrator to visualize data distributions in divergent bar graph formats. DataWrapper and MATLAB were used for additional data visualizations.

Statistics were used in the form of MATLAB *ttest* functions to determine if pre and post-identity differed.

## Results

The assessed course began in the Spring of 2019 and has steadily grown in team size, number of project partners, and number of projects (2D). In the last two years of the course, the enrollment increased by over 400%, with the growth attributed to the increase of project partners.

### Summary of Tech4Wildlife Course Success

During the assessments described in this manuscript, in Spring 2022 and Fall 2022, 18 and 9 students worked on conservation technology projects, respectively. Previously we have reported the overall course objectives[41] as well as themes of the course that have made it successful[30]. The two most prominent that all interviewed students mentioned are two of the foundational framework themes of the VIP Program: interdisciplinary teams and hands-on project experiences[43]. Students have the opportunity to gain multiple semesters of experience in the course. In surveying participants in the course, we utilize the information from the qualitative thematic analysis to quantitatively gauge student identities before and after the Tech4Wildlife VIP course. Students were assessed using the survey displayed in the supplement. From the data provided by the VIP program, since the course’s inception in the Fall of 2019, the course has enrolled a total of 58 students, with 40 students identifying as female, 16 students as male, and two did not mark male or female since its inception.

### Pre-Course Student Identity

Before taking the VIP course, students must fill out an interest form describing their motivation for wanting to join the team. We surveyed the students who had entered the course to gauge what influenced them to participate (**Figure 3**). The most important themes for students applying to the course were: conservation of wildlife (*n* = 5.38 *±* 0.7), interdisciplinary work (*n* = 4.71 *±* 1.03), communication with experts (*n* = 5 *±* 0.93), and working on projects focused on animals (*n* = 5.75 *±* 0.660). Biology and bioengineering typically have higher participation from women than other engineering and science disciplines, and we see similar results here with 70% of the engineering and computer science students in the course’s history identifying as female or non-binary, both of which are underrepresented in the engineering and computer science fields. The least influential portions of joining the class are the sustainable development goals and individual research, as these are likely items that students are unfamiliar with before joining the course.

**Figure 3:**
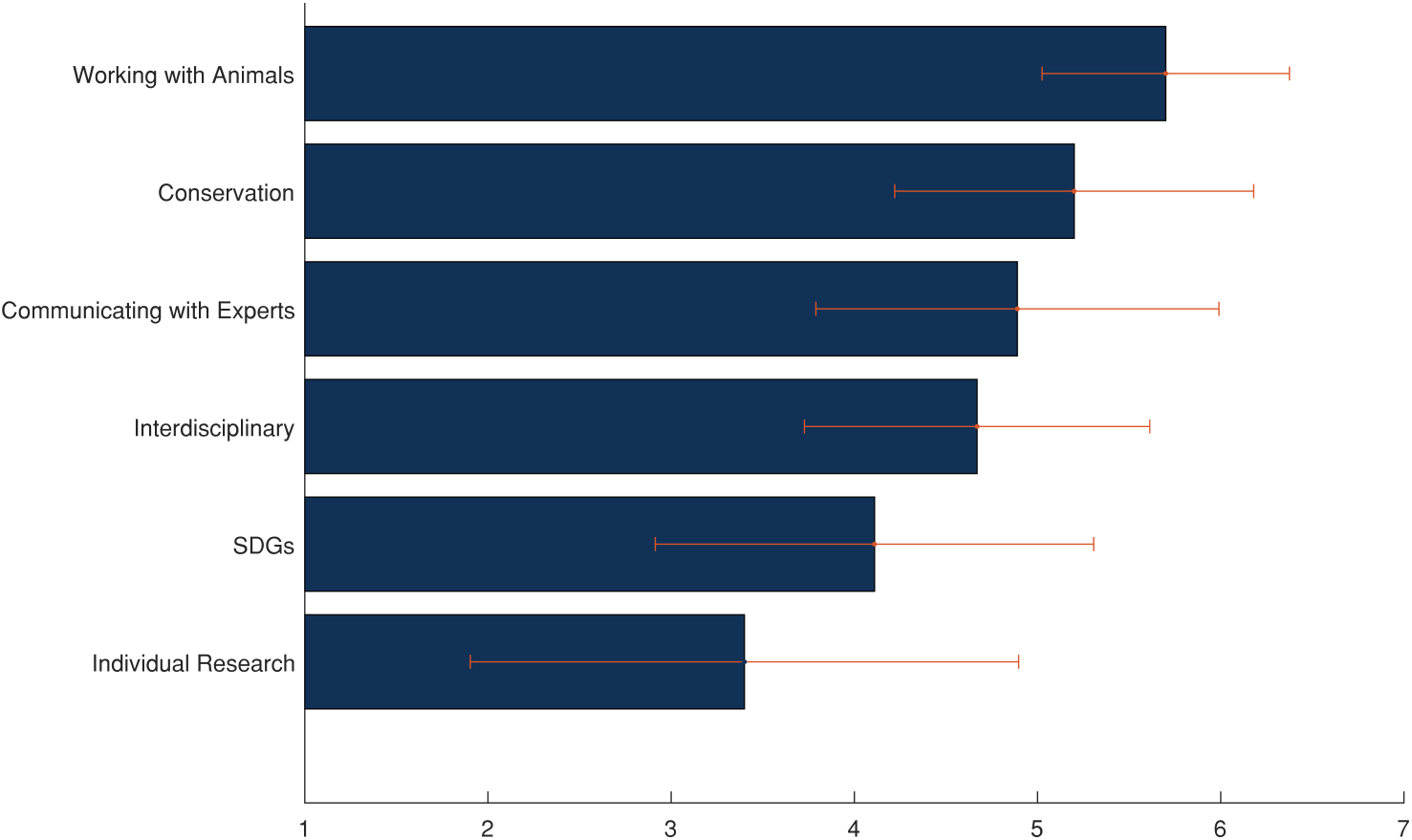
Ranking of the reasoning of students’ reasons for applying to the course ranging from those statements students do not identify with (1) to identify with (7) strongly.

We surveyed a total of 5 engineering students and four computer science students. In analyzing the students’ prior course expectations, we found that most of our respondents were in their first semester in the GaTech4Wildlife VIP course, with 75% indicating it was their first semester. Before attending the course, we gauged students’ identities across the following categories: conservationist, engineer, biologist, technologist, programmer, and environmentalist (**Figure 4**A-C). With such large participation from the engineers and computer scientists, it is logical that students self-identified more strongly in technology-centered identities, including engineer (*n* = 3.85 *±* 1.55), technologist (*n* = 4 *±* 2.07), and programmer (*n* = 3.86 *±* 1.64). Students identified less strongly as biologists (*n* = 1.71 *±* 1.98), conservationists (*n* = 3.1 *±* 1.7), and environmentalists (*n* = 3 *±* 1.58).

**Figure 4:**
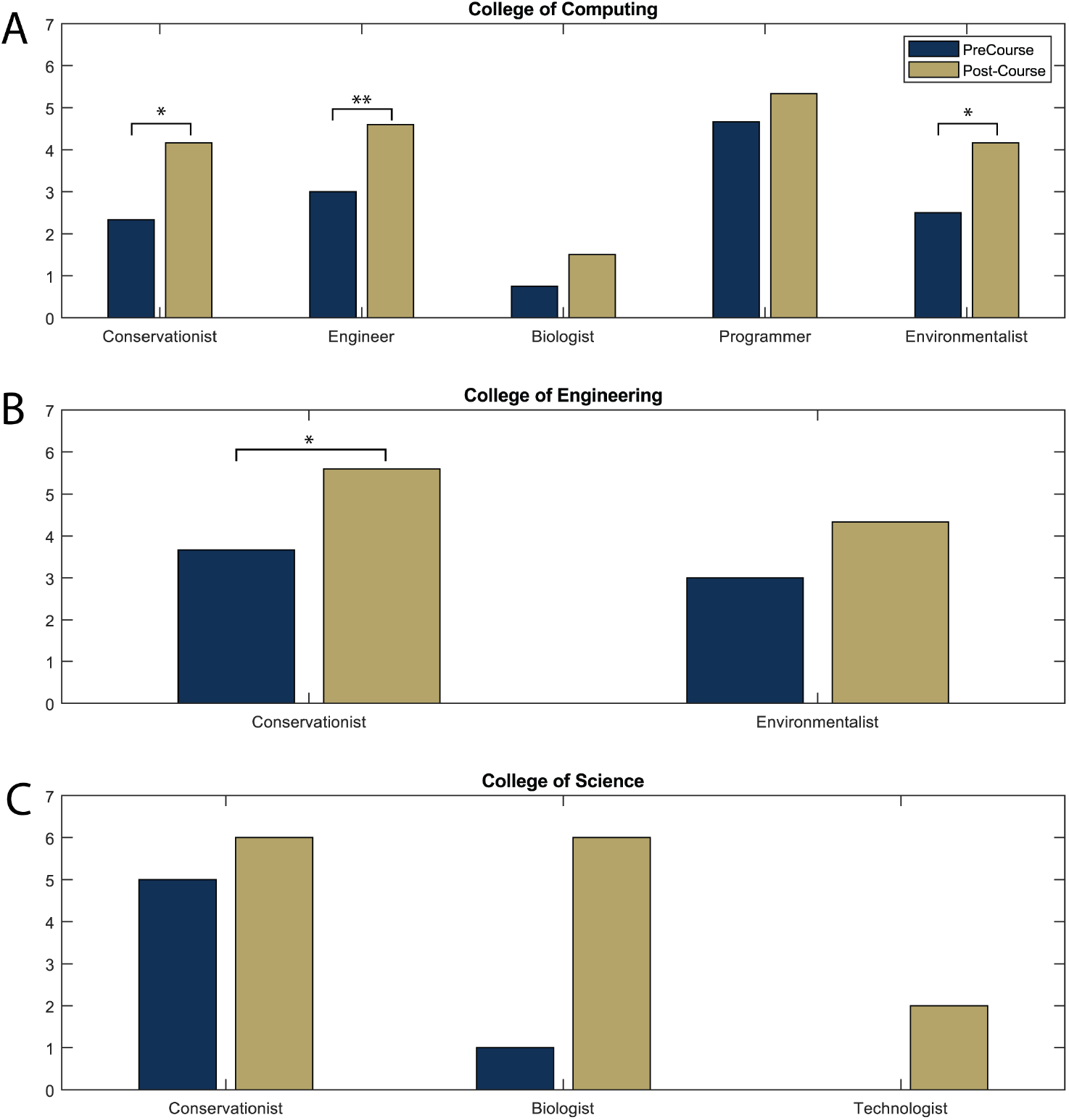
Pre and Post-course identity of students that have participated in the course with all shown cases of increase in students including A) College of Computing student growth of (n=6) students, B) College of Engineering identity growth during the course (n=3) students, and C) College of Science individual growth (n=1) during the course. P-value indicators of * (*p <* 0.05) and ** (*p <* 0.01).

Students in the College of Sciences students had opposite identities with engineers and computer scientists, indicating large differences. These identities likely pair with the courses taken before joining the VIP. We gauged this by indicating the number of courses students had taken before joining. Nearly all engineers and computer scientists (*n* = 85%) indicated they had taken less than two courses on sustainability and environmentalism. In contrast, they indicated more than four courses on engineering design, computer science, and project-based learning. This aligns with Georgia Tech’s project-based curricula, such as ME 2110[46], an introduction to computer science for both engineers and computer scientists. The biology experience of students was only found in the students from the College of Sciences, where engineers and computer scientists had little to no experience. Prior work has told us that Tech4Wildlife is the first biology-related course engineering and computer science students have taken since introduction to biology[45]. For post-graduate plans, students in Tech4Wildlife were primarily interested in graduate school, project management, or non-profit work. They did not often cite conservation work or research as a likely next step (**Figure 5**A).

**Figure 5:**
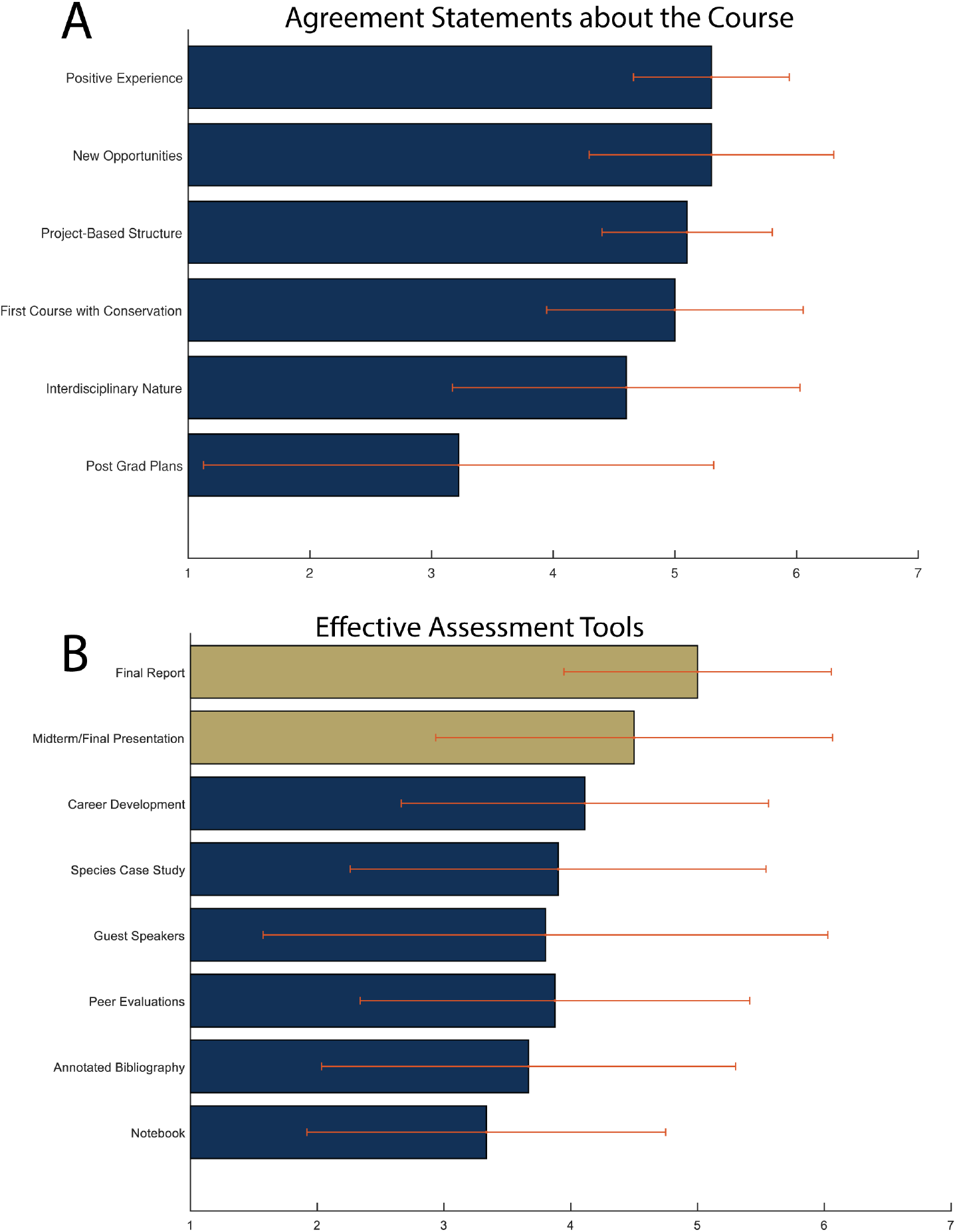
Post course completion evaluation of the course including A) students’ level of agreement from (1) little to (7) greatly agree with statements regarding the overall course. B) Students’ evaluation of which portions of the course are assessment tools, with blue bars indicating formative assessments and gold bars indicating summative assessments.

### Perceptions of Student Assessments

With the class being a project-based course, the only summative assessments are the end-of-the-semester report and presentation. During the semester, several formative assessments have little impact on the student’s overall course grades (**Figure 5**B). When students were surveyed about specific course elements, the two most beneficial on a scale from zero to six were the summative assessments of the final report (*n* = 5.14*±*1.12) and the midterm/final presentations (*n* = 4.5*±*1.73). Students also enjoyed the career development aspect of the course near the end of the education phase module (*n* = 4.3 *±* 1.58). The least beneficial experiences, as reported by the students, were the annotated bibliography (*n* = 3.4 *±* 1.73) and the notebook (*n* = 3 *±* 1.41). Every other assessment/portion of the course: guest speakers, peer evaluations, and specific case studies all exhibited means above 3.5, indicating that students felt they were helpful in the experience but were not the primary factors.

### Applications of Themes to Engineering Curriculum

We surveyed students who had completed the course about the themes found during detailed one-on-one student interviews. The primary course themes, as per described in detail in Schulz et al. for the GaTech4Wildlife VIP course, were as follows: [45]

- Hands-on team project experiences in conservation tech,
- Exposure to research opportunities for individuals to apply learned knowledge to conservation,
- Eye-opening perspective on using technology to conserve wildlife,
- Connections made among varying disciplines during the course.

Each of these respective themes was described by over 75% of students interviewed. These questions included discussions of exposure and the impact on post-graduate plans. Overall we found that the statements were the most agreed upon were students felt that the course positively impacted their experience at Georgia Tech (5.38 *±* 0.7), the project-based structure contributes to the effectiveness (5.38*±* 0.7), the class contributes to conservation effectiveness, (5.38*±* 0.7), and the course was their first experience with conservation, (4.38 *±* 0.7). The only statement that students disagreed with is that the course has influenced their post-graduate plans (3 *±* 2.27). This question had the most significant variance of almost any question on the survey, likely indicating that for some students, it had a very high impact, but for others, it had almost none. This could be because many students were exposed to new opportunities. Still, with the conservation technology field being so new and novel, there are currently very few post-graduate jobs in the field.

### Post Course Completion Identity

We asked students about their post-course completion identities after their first semester of the VIP course. All student participants indicated increases in all identities except programmers. We found increases in student identities of conservationists, engineers, biologists, technologists, and environmentalists. In performing a two-tailed t-test of statistical significance, we found significant growth in the identity of all participants in the identities of environmentalists and conservationists (**Figure 4**A-C).

When we separate the College of Computing students and the College of Engineering students, we see particular identity changes. For the computer science students, engineer and technologist identities increased while maintaining a constant programmer identity. The computer scientists also nearly tripled their identity as conservationists (5 *±* 1) and environmentalists (4.25 *±* 1.4), which may be because no computer scientist indicated prior courses on sustainability. In the College of Engineering, we see similar identity increases for conservationists (5.2 *±* 1) and environmentalists (4.7 *±* 1). Still, engineers also exhibited an increase in identity as biologists (2.7 *±* 0.6), up from (1.2 *±* 1) before the course, possibly due to the trans-disciplinary nature of the course’s project-based goals and human-wildlife-centered framework. Students in the biological sciences did not indicate increased identity as programmers. Still, their identity as engineers and technologists increased, allowing College of Science students to gain engineering education outcomes through the course. Significant identity changes (*p <* 0.05) occurred with both computing and engineering students identifying stronger as conservationists after the course and college of computing students identifying more robust as environmentalists and engineers.

### Student-Initiated Expansion to Student Organization

After the course’s early successes with project completions at local conservancies, including Atlanta Botanical Gardens, Zoo Atlanta, and Atlanta Coyote Project, there began to be too many students applying for the course to allow all to participate. Based on our experience, if more than 25 students take the course, there are too many teams or too few teams with too many people, making it harder to maintain engagement. In peer evaluations with class sizes of 30, several students began to feel that they were not contributing adequately to their projects. With the increase in members that joined the class the course evolved into a student organization as well as the course. The student organization Tech4Wildlife at Georgia Tech hosts a yearly Hack-a-Thon on sustainability, has guest speakers, and works on local outreach on various events. The student organization officially started during the Spring of 2022 and remains active with recruiting new members and expanding to new events.

## Discussion

### Addressing the Sustainability Gap

With the recent adoption of sustainable development goals, universities must adopt new and novel fields, such as conservation technology, into their curriculum. Several computer science programs nationwide have adopted a similar field, AI4Good, based on Microsoft AI4Good and Google Earth Initiatives[47]. Programs, including the Vertically Integrated Projects program, study abroad, and senior design, give students an excellent platform to practice their sustainable engineering practices. To address this sustainability gap, universities need to work on partnering with not just industry partners but also non-profits and sustainability organizations such as Engineering for One Planet[48], zoological organizations[30], or sustainability startups to engage in the next generation of sustainability outcomes. Partnering with these organizations and change-makers will allow universities to continue their rigorous engineering outcomes but focus the engineering design process on challenges proposed by sustainability-focused organizations. A prime example is the partnership between the Boeing corporation’s EcoDemonstrator program. The EcoDemonstrator program is a testing plan that partners with the University of Washington to allow students to engage in sustainability-minded testing in real-world industry applications [49].

### Bio-mimicry Related to Sustainability

There have been recent advances in developing Bio-Inspired and Biomimicry-based courses and modules to enhance engineering education. The field of bio-inspired design (BID) has had success in being taught in multi-disciplinary and engineering classes at the high school, undergraduate, and graduate curriculum levels [50]. There are numerous reasons for the proliferation of such courses: spurring and teaching innovation practices, the focus on sustainability and the SDGs, and recruiting more women into engineering classes where they are typically underrepresented. Biology and Bio-engineering typically boast 50-60 % participation rates by women students, while participation more broadly in engineering still hovers around 20%[51]. It has been hypothesized that incorporating biological content into engineering design courses may improve participation from women students. However, teaching BID is not without its challenges.

BID requires deep knowledge of both biological structures and engineered systems. This creates a challenge for both teachers and students to be sufficiently versed in both spaces to actually apply BID appropriately, as opposed to naively[52, 33]. In efforts to reduce the need for multi-disciplinary knowledge, websites such as Ask Nature and even AI frameworks for accessing biological organisms have been proposed; however, this approach can be exploitative and diminish the contributions of biologists serving on design teams. Finally, BID does not explicitly address sustainability; BID can be perceived as harvesting solutions from biology to serve human-engineered systems without taking care to preserve biology or understand biology for sustainability purposes. While some BID curricula may also include sustainability elements, it is possible that BID curricula cultivate the view that biology is most beneficial in service to engineering.

Conservation Tech may offer a pathway that engages women and diverse students based on its relationship with biology and its altruistic nature with fewer risks of exploiting biology or biologists. This is not to say that BID should not be taught, but instead that CT may pose a greener pathway for broader engagement in the SDGs while the focus of BID is on pushing the boundaries of engineered systems and not explicitly on conservation.

### Course Challenges

A significant challenge with teaching a course on engineering and computer science’s influence on conservation science is addressing misconceptions. Students can often have misconceptions about conservation through ideas such as scooping plastic out of the ocean, hunting, fishing, trophy hunting, zoo benefits for conservation, de-extinction, and more. As this is such a significant challenge, we began each class by addressing one of the misconceptions in the conservation sector by watching a video and having a think-pair-share discussion openly between different people. Pairs would be organized to support interdisciplinary conversations to help discussions become more holistic, and prompting questions were given to help generate student ideas.

Additional challenges in implementing a course like this are the politicization of conservation and sustainability, particularly in the United States. With ideas such as climate change and sustainable energy, green engineering, and more becoming politicized, we grounded much of our discussions in the class with Sustainable Development Goals.

### Starting Your Own Conservation Tech Course

For beginning a course on the idea of sustainability, we highly recommend that instructors look to their research and interest and begin with incorporating sustainability practices into their lessons. Taking something like a Civil or Environmental Engineering course and utilizing novel hardware to include sustainability through the lens of one of the Sustainable Development Goals will significantly help the success of the course moving forward. It is also possible that This course was constructed based on the author’s previous experiences with biologically-inspired design education and conservation science. The idea of interdisciplinary education combined with conservation science started this course. As instructors look to teach conservation technology and conservation tool creation to engineers, it is essential to find community conservation partners to help engage. In our class, we worked closely alongside non-profits and organizations like Zoos, which helped foster a collaborative research agreement[30].

We highly recommend partnerships with conservation organizations as this can allow guest speakers and future project ideas to be created in a collaborative space. A significant challenge in the conservation sector is the lack of funding and resources. Providing students with a project working with a local organization will also allow for potential fieldwork and longer-term projects as the community is invested in the long-term success of the collaboration.

## Conclusion

In this paper, we demonstrate the conservation technology taught in a project-based setting allows for the interdisciplinary growth of engineers and computer scientists to grow in their sustainability mindset as well as their identities as sustainable, conscious engineers. Additionally, we see the growth of biologists in the hardware and software identities through interdisciplinary and long-term projects that engage conservation organizations. Finally, we see the potential for this type of field to successfully be implemented into engineering curricula as having long-term outcomes of identity for post-graduate years and provide specific advice for pre-course, course-execution, and course improvement and challenges for others that would like to begin a project-based course on conservation technology.

## Acknowledgements

We would like to acknowledge the Serve-Learn-Sustain program at Georgia Tech, including the Sustainability Faculty Fellows program, which supported this work, the Georgia Tech Vertically Integrated Project (VIP) program, and finally, the Georgia Tech Research Institute (GTRI’s) support of this VIP course, especially to Robert Wallace and Paula Gomez. Additionally, we thank David Hu, Ed Coyle, Kate Williams, Tammy McCoy, and Amanda Lai who were involved in the development of the beginning of the Tech4Wildlife course.

